# Suidae iPSC-derived macrophages as models for investigating susceptibility and resilience to African swine fever virus

**DOI:** 10.64898/2026.04.22.719209

**Authors:** Tom M. Watson, Lynnette C. Goatley, Stephen Meek, Lel Eory, Sylvia Kohler, Nicholas Berkley, Sofia Sternberg, Melany Jackson, Amy Findlay, Isabella Hoskins, Simon Girling, Joseph Mee, Alan L. Archibald, Finn Grey, Falko Steinbach, Helen Crooke, Christopher L. Netherton, Tom Burdon

## Abstract

African swine fever virus (ASFV) causes a lethal haemorrhagic fever in pigs and spread of this disease threatens many pig species (Suidae) globally. By contrast, ASFV infections in the natural evolved hosts, the warthog and bushpig, are subclinical. The macrophage (Mφ) is the primary target of ASFV and species-dependent responses in Mφs are presumed to influence disease susceptibility. In an attempt to model these differences *in vitro*, we generated transgene-regulated induced pluripotent stem cells (iPSCs) from domestic pig, wild boar, red river hog and warthog, and confirmed that their corresponding iPSC-derived Mφs (iPSCdMs) supported infection and replication of ASFV. In contrast to the other species, however, warthog iPSCdMs did not induce interferon α upon infection by either virulent or attenuated ASFV. iPSCdMs may therefore represent an experimental system to understand how ASFV infection of Mφs contributes to disease and aid development of strategies to combat this economically and environmentally devastating pathogen.

## Introduction

ASFV is a large, ≥200 kb double stranded DNA virus of the Asfarviridae family that is present in a sylvatic cycle between warthogs (*Phacochoerus africanus*) and soft ticks (*Ornithodoros* spp.), as well as in bushpigs (*Potamochoerus larvatus*) in Sub-Saharan Africa (Galindo & Alonso, 2017; Penrith & Kivaria, 2022). Despite harbouring significant levels of virus these wild porcine species do not manifest clinical symptoms following infection (Anderson et al., 1998; Blome et al., 2013; Oura et al., 1998; G. R. Thomson et al., 1980). By contrast, ASFV infection of domestic pigs (*Sus scrofa domesticus*) and wild boar (*Sus scrofa*) is lethal, causing a haemorrhagic fever that kills most animals in less than 10 days (Galindo & Alonso, 2017; Gómez-Villamandos et al., 2013; Sánchez-Cordón et al., 2018). At present there is no broadly effective, available vaccine against ASFV and attempts to control spread of the virus have been limited to large scale culling of infected and neighbouring herds (Netherton, 2021; Viltrop et al., 2021; Wang et al., 2023). The spread of ASFV from infected farmed and wild porcine populations poses a continual threat to pig production. ASFV outbreaks in Central and Eastern Europe, Asia, and most recently in Spain have had devastating effects on commercial pig farming (Sánchez-Cordón et al., 2018). The latest introduction of ASFV in China in 2018 caused national shortages of pork meat and estimated production losses of > $111 billion (You et al., 2021). In addition to the welfare issues associated with controlling ASFV on farms, the spread of the virus threatens to destroy many rare indigenous wild porcine species, most imminently in South East Asia (Cowled et al., 2022; Daniel et al., 2025). There is a pressing need therefore to develop new strategies to combat the effects of this devastating pig pathogen (Wang et al., 2023).

ASFV infects cells of the mononuclear phagocyte system, principally the Mφs, which are key effectors of the innate immune response and regulators of the adaptive immune system (Gómez-Villamandos et al., 2013). Virus entry into Mφs is likely complex, exploiting multiple redundant entry mechanisms including receptor/clathrin-mediated endocytosis, micropinocytosis and apoptotic mimicry (Hooper et al., 2024). Nevertheless, genome-wide CRISPR mutagenesis screens in a wild boar cell line have pointed to the involvement of the SLA-DM MHC class II and TMEM239 proteins in early virus infection (Pannhorst et al., 2023; Shen et al., 2024). Following entry, ASFV is trafficked to the late endosomes where the viral core enters the cytoplasm and proceeds to replicate in perinuclear viral factories (Hernáez et al., 2016). New virus particles bud out through the plasma membrane and intracellular virus, that is also infectious, is released after host cell lysis (Andrés et al., 2001). The ASFV genome encodes more than 200 proteins (Cackett et al., 2022), some of which are involved in modulating host responses including suppressing the activation of a type I interferon (IFN) response and regulating cell survival through programmed cell death pathways (Dixon et al., 2017, 2019; Netherton et al., 2023; Reis et al., 2023; Wu et al., 2021). After the initial phase of infection an excessive release of pro-inflammatory cytokines likely contributes to a lethal widespread haemorrhage in domestic pigs (Franzoni et al., 2023). Despite carrying high viral titres following ASFV infection, warthogs and bushpigs do not exhibit significant clinical signs (Anderson et al., 1998; Oura et al., 1998; Thomson et al., 1980). By contrast, infection of domestic pigs and other vulnerable porcine species leads to the overreaction of a maladapted immune response responsible for the damaging pathology.

ASFV persists predominantly in eastern and southern African in a sylvatic cycle between *Ornithodoros* soft ticks and common warthogs *Phacochoerus africanus*. Whilst, newborn warthogs in particular exhibit high, sustained, levels of virus deemed sufficient to infect feeding ticks present in the warthog burrows, adult warthogs, by contrast, show lower, more transient levels of viraemia (Auer et al., 2025). In experimental infections of warthog and bushpigs, high levels of virus are sustained for less than a month before declining to below the level of detection by 4 months (Anderson et al., 1998; Oura et al., 1998; Thomson et al., 1980). *In vitro* infections of warthog, bushpig and pig *ex vivo* cultured leukocytes showed similar patterns of ASFV growth over a 7-day period, consistent with the rapid increase in viraemia observed during the acute phase of infection in all these species. Crucially, however, there have been no reports of infected warthogs or bushpigs exhibiting clinical signs of disease, indicating that these suids have a robust, evolved resilience to ASFV.

Comparing the response of domestic pigs and African wild pigs to ASFV infection can provide important insights into mechanisms that regulate susceptibility and resilience to ASFV. However, experimental access to wild or captive animals and performing reproducible *in vivo* and *in vitro* challenges is both logistically and ethically challenging. Pluripotent stem cells (PSCs) may provide an alternative, genetically tractable and limitless source of differentiated cells that can be used to model host-virus interactions (Rajab et al., 2018). We have previously shown that PSCs derived from pig pre-implantation embryos serve as a source of *in vitro* differentiated pig Mφs that exhibit many key Mφ characteristics including their susceptibility to the pig Mφ-specific viruses, Porcine reproductive and respiratory syndrome virus (PRRSV) and ASFV (Meek et al., 2022). However, due to extremely limited availability of suitable embryos, reliable derivation of embryo-derived PSCs from wild animals is not practicable. By contrast, conversion of somatic cells into a PSC-like state using iPSC reprogramming is feasible using cells derived from biopsied tissue (Hutchinson et al., 2024; Takahashi & Yamanaka, 2016). In this report we describe the derivation of iPSC lines from domestic pig and wild porcine species and their differentiation into Mφs as a biologically relevant cell culture model for studying host-ASFV interactions.

## Results

### Derivation of pig iPSC lines

We used a modification of the reprogramming protocol first described by Gao and co-workers, to generate pig iPSC primary colonies (Gao et al., 2019). Pig embryonic fibroblasts were electroporated with three piggyBac-based vectors that direct doxycycline-regulated expression of standard reprogramming pig factors OCT4, SOX2, KLF4 and MYC (OSKM); pig NANOG and human LIN28B (NL), RARG and LRH1 (RL). Electroporated pig fibroblasts were plated on gelatin coated dishes in M15 medium (GMEM + 15% FCS, Vitamin C, human LIF) supplemented with 1 mM sodium butyrate (NaB) and 1 μg/mL doxycycline (M15G+Dox medium). After 7 days the cells were transferred to plates containing irradiated STO fibroblast feeder cells and cultured in M15G+Dox medium. Loose clusters of small cells first emerged between 7-10 days, and some progressed to form colonies with a distinctive compact stem cell morphology identifiable at 14-28 days (Figure 1A). Colonies displaying stem cell morphology were picked and expanded on irradiated STO feeder cells in M15G+Dox medium to establish cell lines that expressed the stem markers alkaline phosphatase and EPCAM (Figure 1B-D). Basic multilineage differentiation potential of lines was assessed by their capacity to generate morphologically distinct cell types following doxycycline withdrawal and embryoid body differentiation. Based on this criteria, two pig iPSC lines, piPSC1 and piPSC2, that displayed diverse differentiation potential were selected for further experiments.

**Figure 1.**
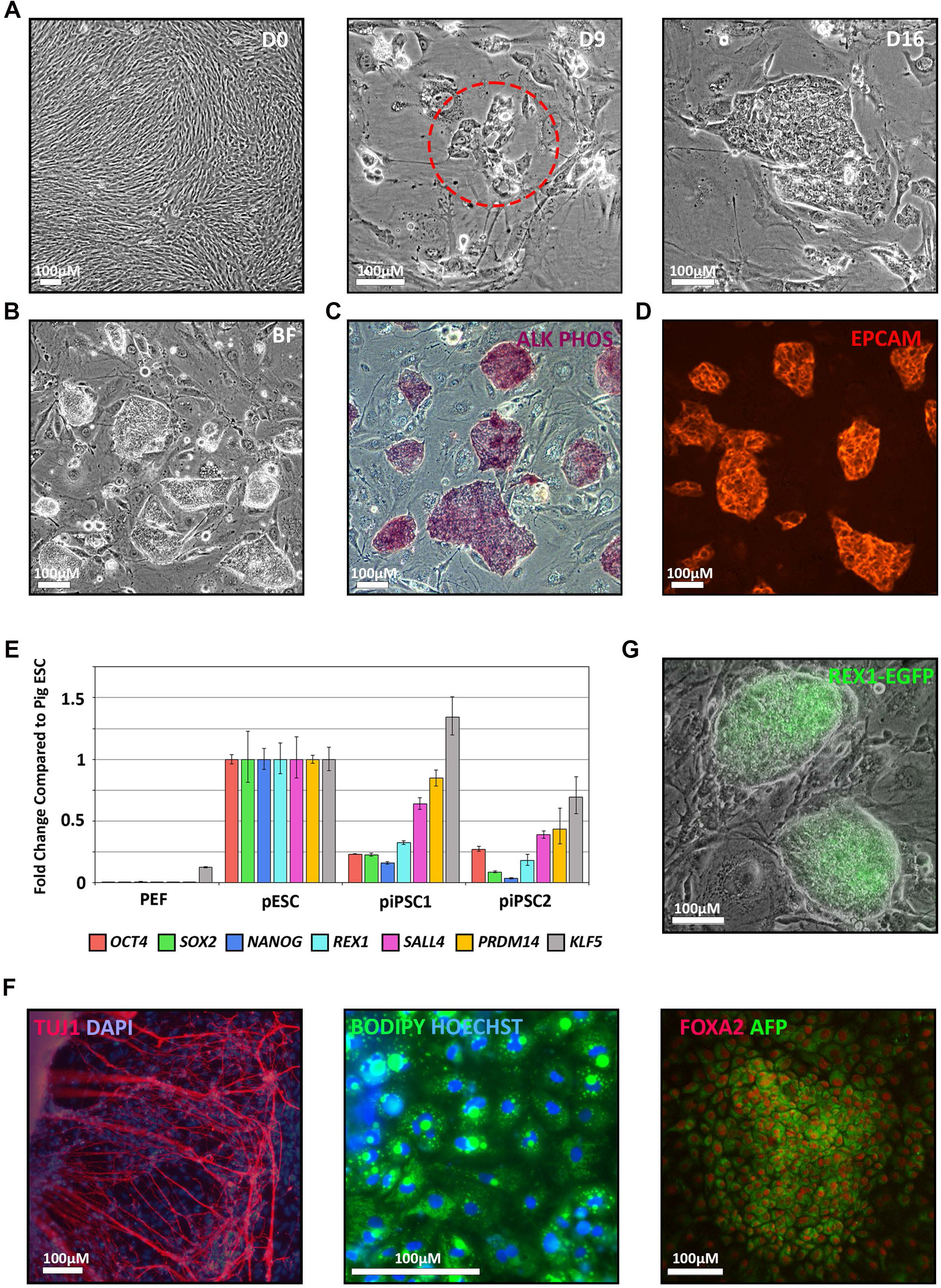
Generation and functional characterisation of domestic pig iPSCs. (A) Bright field images showing the time course (Days 0,9,16) of iPSC colony formation during reprogramming of pig embryonic fibroblasts (PEF). (B) Representative bright field image of pig iPSC1 (piPSC1) after expansion. (C) Alkaline Phosphatase staining of piPSC1. (D) Fluorescent image of Immunocytochemical (ICC) staining for EPCAM (Red) expression in piPSC1. (E) RT-qPCR data showing expression of endogenous pluripotency genes in piPSC1 and piPSC2 compared to porcine ESCs (pESCs) and fibroblasts. Mean ±SD of 3 independent experiments. (F) Fluorescent images of piPSC-derived ectoderm (beta3-Tubulin: TUJ1 stained neurons), mesoderm (BODIPY stained adipocytes) and endoderm (AFP and FOXA2 stained embryonic endoderm). (G) Merged bright field and fluorescent images of a piPSC1 *REX1*-*EGFP* knock-in clone.

To assess the extent of reprogramming of the pig iPSCs, we compared transcript levels of the endogenous pig pluripotency genes *OCT4, SOX2, NANOG, REX1, SALL4, PRDM14*, and *KLF5* with that of an established pig embryo-derived EPSC line (Gao et al., 2019) (Figure 1E). Expression of the endogenous genes, with the exception of KLF5, were all lower in both iPSC lines. Nonetheless, differentiated piPSC1 cultures produced beta3-tubulin (TuJ1) expressing neurons, lipid containing immature adipocytes and AFP+FOXA2 positive hepatocyte-like cells, demonstrating that the iPSCs were capable of generating cells of all three germ layers and that transgene expression had conferred a pluripotent stem cell-like state (Figure 1F, S1).

To further confirm pluripotency and establish gene editing potential of the pig iPSC lines, we electroporated the piPSC1 cells with a pig *REX1-EGFP-IRES*-puro stem cell-reporter vector and Cas9/Rex1gRNA RNP complexes and selected puromycin resistant colonies as described previously (Meek et al., 2022). PCR analysis of genomic DNA from expanded clones confirmed that the EGFP reporter was correctly inserted at the 3’ end of the endogenous pig *REX1* stem cell gene in most clones, and EGFP expression was detected in the majority of the morphologically undifferentiated stem cells consistent with acquisition of a pluripotent state (Figure 1G and S1).

### Generation and molecular characterisation of pig iPSC-derived Mφs

We next differentiated the iPSCs using a three-phase Mφ differentiation protocol previously applied to pig PSCs (Meek et al., 2022). Mesoderm and early haematopoietic differentiation was induced through embryoid body (EB) suspension culture in the presence of BMP4, SCF, and VEGF cytokines. EBs were then allowed to attach to form a monolayer and induced with IL-3 and CSF1 to produce myeloid derivatives (Figure 2A). After 10-14 days in this second phase, floating aggregates and scattered attached cells with Mφ-like morphology became evident. In the following 3 weeks, the production of the floating cells increased and when harvested from the culture medium these cells could be converted in the third phase in medium containing CSF-1 into cells with mature Mφ-like morphology (Figure 2A).

**Figure 2.**
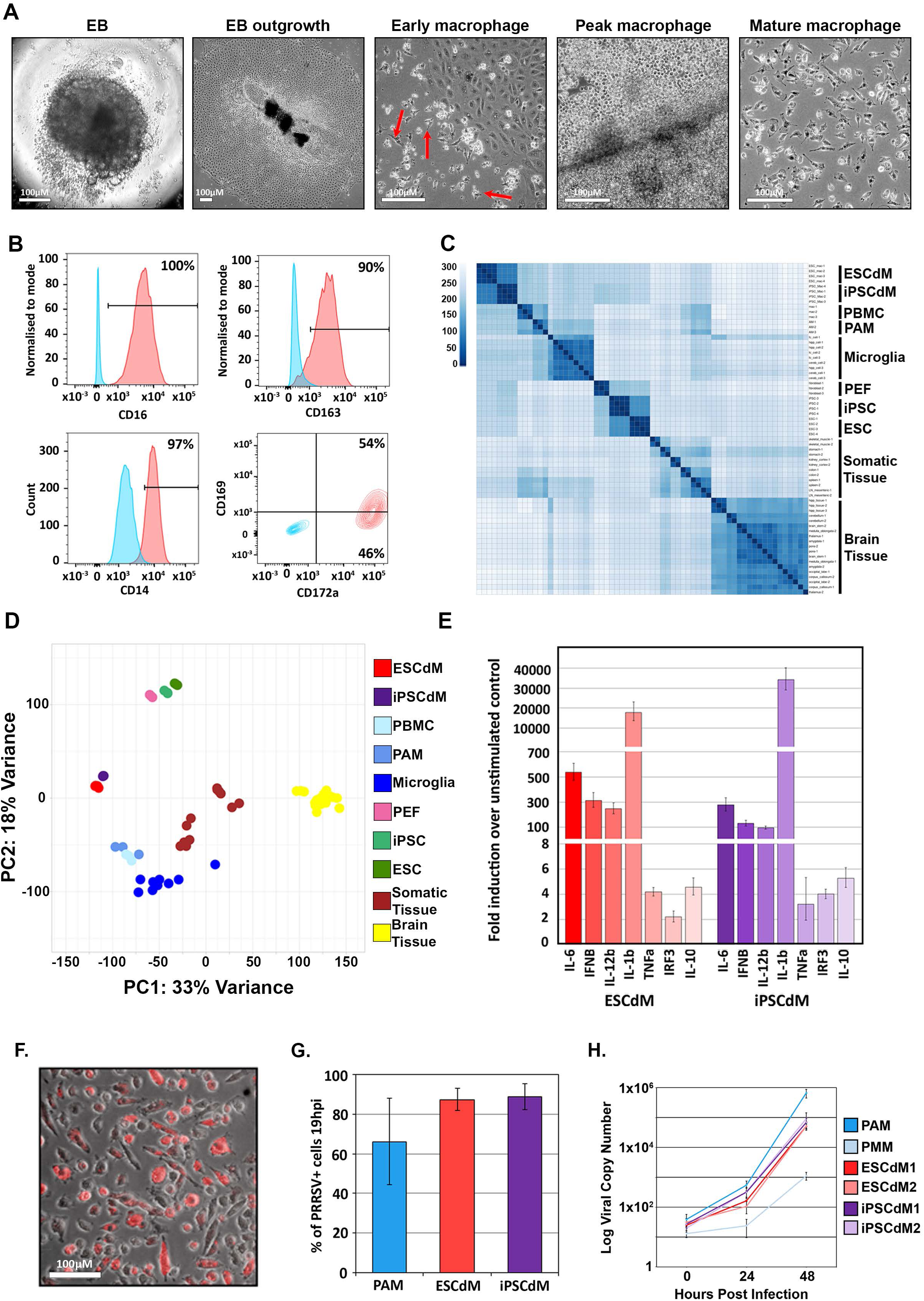
Characterisation of domestic pig iPSCdMs. (A) Representative bright field images of different stages of macrophage (Mφ) differentiation. Red arrows identify early iPSCdMs. (B) Flow cytometry analysis of iPSCdMs stained for Mφ surface markers (Red) relative to isotype controls (Blue). (C) Heatmap showing sample similarity based on the Euclidean distances. Darker colour represents closer relationships between samples based on their expression profiles. (D) PCA plot of the first two principal components (PC1 and PC2) for tissue-specific gene expression. (E) RT-qPCR analysis of immune response in ESCdMs and iPSCdMs following 6 hrs induction with 200 ng/mL LPS. Mean ±SD of three independent experiments. (F) Composite bright-field and fluorescent image showing phagocytosed pHrodo-Red beads in pig iPSCdMs at 3 hrs post addition. (G) Graph showing the proportion of Mφs stained for PRRSV nucleocapsid protein at 19 hours post infection (hpi). Mean ±SD of three independent experiments. (H) qPCR analysis of ASFV genome copies in supernatants from 24 hpi and 48 hpi Mφs. Mean ±SD of 4 technical replicates.

To evaluate the molecular profile of the iPSCdMs we examined their cell surface protein expression and transcriptional profile. Flow cytometry of mature (phase 3) iPSCdMs stained using fluorophore conjugated antibodies showed that most cells in the cultures expressed characteristic Mφ proteins CD16, CD14, CD172a, and CD163 and approximately half expressed modest levels of CD169 in a pattern resembling that of embryonic pluripotent stem cell-derived Mφs (PSCdMs) and *ex vivo* Mφs (Meek et al., 2022) (Figure 2B, S2). iPSCdMs expressed characteristic Mφ transcripts, albeit at reduced levels compared with pig alveolar Mφs (PAMs) maybe reflecting a relatively immature state. By contrast, expression of the reprogramming transgenes was not detected in iPSCdMs (Figure S2). To assess the global gene expression profile of iPSCdMs we compared their transcriptome with that of PSC-derived Mφs, ex-vivo pig Mφs, iPSCs, ESCs and pig tissue samples. Heatmap and hierarchical clustering analysis of the 100 most highly expressed genes in these samples showed that the transcriptional profile of the iPSCdMs most closely resemble the embryo-derived PSCdMs, and these samples in turn are similar to *ex vivo* peripheral blood monocyte cells (PBMC)-derived Mφs and PAMs, and then more distantly related to microglia (Figure 2C, S2). Multidimensional scaling at a genome-wide level confirmed the relationship between the two types of stem cell-derived Mφs and their proximity to the other *ex-vivo* Mφs, but divergence from their respective parental stem cells and other pig cell types (Figure 2D). In conclusion, the analyses of cell surface protein expression and gene transcription profiling strongly indicate that iPSCdMs closely resemble embryo-derived PSCdMs.

### Functional characterisation of iPSCdMs

To evaluate iPSCdMs functionality we measured the phagocytic activity of iPSCdMs by monitoring uptake of pHrodo zymosan beads that fluoresce in the acidic environment of the phagosome. Within 3 hours of addition of the beads to the culture medium the majority of the iPSCdMs contained fluorescent beads in line with high levels of phagocytic activity previously observed for ESCdMs (Meek et al., 2022) (Figure 2F). To test the response of iPSCdMs to immune activators, iPSCdMs, and ESCdM controls were treated with the TLR4 ligand lipopolysaccharide (LPS) and downstream gene activation was measured by RT-qPCR (Figure 2E). Similar patterns of LPS-mediated increases in mRNAs for cytokines IL6, IFNβ, IL-12β, IL-1β, TNFα, IL-10, and transcription factor IRF3 were obtained for iPSCdMs and ESCdMs. We also assessed infection of iPSCdMs by two pig Mφ-specific viral pathogens, PRRSV and ASFV. Approximately twenty hours after incubation with PRRSV, >90% of iPSCdMs expressed PRRSV p63 nucleocapsid protein as determined by flow cytometry and immunocytochemistry, at levels equivalent to that supported by ESCdMs and PAMs (Figure 2G). Similarly, ASFV DNA was detected by qPCR in culture supernatants at 24 and 48 hrs after infection of iPSCdMs at levels equal to ESCdMs, and comparable to *ex vivo* pig Mφs (Figure 2H). Collectively, based on the molecular and functional characteristics tested, iPSCdMs are equivalent to ESCdMs and may represent a useful alternative *in vitro* model for studying Mφ-pathogen interactions in pigs.

### Generation of iPSC and iPSCdMs from wild suids

The equivalency of pig iPSCdMs and ESCdMs suggested that transgene-dependent iPSCs might also prove useful in deriving PSCs from other Suidae species, particularly in circumstances where embryos are unavailable. Of particular interest were species that differ from domestic pigs in their susceptibility to ASFV, such as common warthog (*Phacochoerus africanus*) and red river hog (*Potamochoerus porcus*), a close relative of the bush pig (*Potamochoerus larvatus*) (Auer et al., 2025; Balboa et al., 2024). We therefore applied the transgene-dependent reprogramming protocol to primary fibroblasts obtained from warthog, red river hog and wild boar. At 14-28 days post-transfection with piggyBac reprogramming vectors, stem cell colonies were picked and expanded to generate cell lines for red river hog and wild boar. However, in standard M15G medium the predominant morphology of warthog colonies was irregular and incoherent, with many loosely attached cells. Increasing the concentration of sodium butyrate improved the morphology but inhibited growth. On the basis that growth inhibition might be an off-target effect associated with higher concentrations of sodium butyrate, we added another HDAC inhibitor valproic acid to M15G+Dox medium and found that this improved the morphology of the warthog iPSC colonies without inhibiting growth. Expanded iPSC clones were subjected to a scaled-down Mφ differentiation protocol to identify useful differentiation-competent iPSCs. Clones that produced embryoid body outgrowths containing rapidly growing epithelial-like cells and signs of Mφ production were selected for further evaluation. Differentiation-competent lines were generated for each species (Figure 3A). However, the efficiency varied between lines and in some cases only a minority of the EB outgrowths produced Mφs. Reduction or removal of HDAC inhibitors for at least a passage before initiating the differentiation protocol improved efficiencies. We considered that heterogeneity of the extent of reprogramming within clones might be responsible for some of the variation and that a second round of reprogramming driven by reactivation of the reprogramming transgenes in iPSCdMs might select a subpopulation of differentiation-competent iPSCs. To test this idea iPSCdMs from pig, red river hog and wild boar iPSC lines were plated on STO feeder cells in M15G + Dox medium to induce re-expression of the reprogramming factors. Undifferentiated iPSC-like clones emerged from the Mφ cultures after 10 days and were then re-tested for their capacity for Mφ differentiation (Figure S3). In comparison with the parental clone, each of the secondary reprogrammed iPSC lines exhibited a marked (∼30%) increase in the EBs producing Mφs, representing a significant improvement in the utility of the iPSCs (Figure S3). Indeed, wild boar, red river hog and warthog iPSCs differentiated into immature liver-like cells and neurons, representing endoderm and ectoderm germ layer derivatives, respectively (Figure S4). In addition, CRISPR-mediated targeted insertion of the EGFP fluorescent reporter gene at the 3’ end of the *REX1* gene in red river hog iPSCs, as described previously for pig iPSCs, confirmed that wild pig iPSCs were also amenable to standard gene editing techniques (Figure S4).

**Figure 3.**
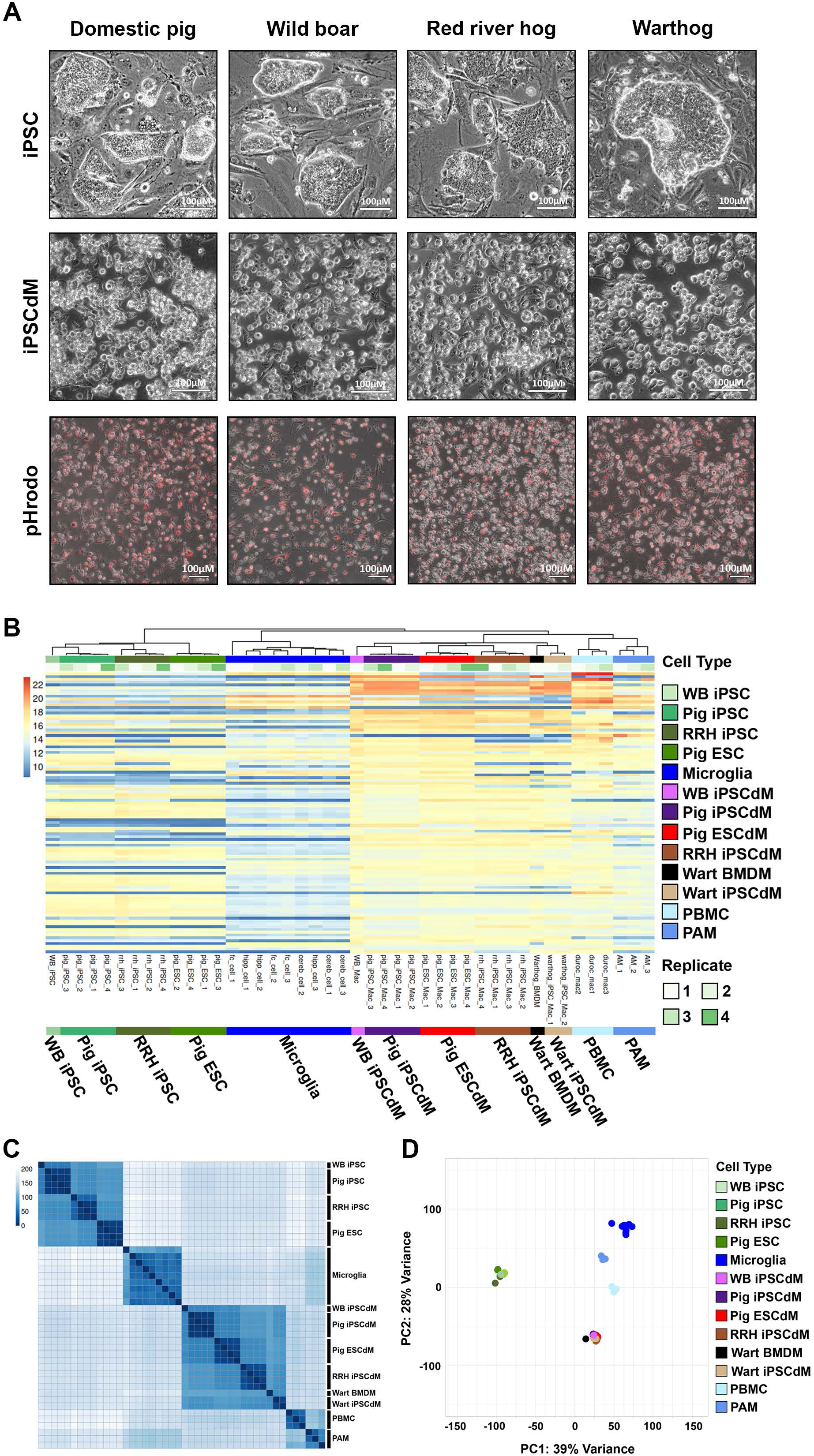
Generation of iPSCdMs from wild suids. (A) Representative bright field and fluorescent images of suid iPSC clones, iPSCdMs and phagocytosis of pHrodo-Red beads by iPSCdMs at 3 hrs post addition. (B) Heatmap of the hundred most highly expressed genes in suid PSCdMs and *ex vivo* Mφs. Tree indicating hierarchical clustering between samples at top. (C) Heatmap showing sample similarity based on Euclidean distances. Darker colour represents closer relationships between samples based on their expression profiles. (D) PCA plot showing the first two principal components (PC1 and PC2) in cell-specific gene expression.

Application of the Mφ differentiation protocol to selected wild boar, red river hog and warthog iPSC lines produced cells with morphology comparable to pig iPSCdMs that were actively phagocytic, as demonstrated by their efficient internalisation of fluorescent zymosan pHrodo beads (Figure 3A). To assess the broader Mφ identity of the iPSCdMs, we compared their transcriptional profiles with that of pig Mφs and *ex vivo* warthog bone marrow-derived Mφs (Figure 3B, C, D). Heatmap and hierarchical clustering analysis of the 100 most highly expressed genes illustrated that the transcriptional profiles of the iPSCdMs of all the species and pig PSCdM were closely related, collectively similar to *ex vivo* Mφs from pig and warthog and, more distantly, microglia (Figure 3B, C). Moreover, multidimensional scaling at a genome-wide level provided a global view of the similarities between cell and tissue types and confirmed the close relationship between the iPSCdMs and proximity to the other ex-vivo Mφs, but divergence from suid PSCs (Figure 3D).

Warthogs and bushpigs can be infected by ASFV, without succumbing to disease (Oura et al., 1998; G. R. Thomson et al., 1980). To determine if *in vitro*-derived Mφs from warthogs and red river hogs are permissive for ASFV, we infected iPSCdMs from these species with Georgia 2007/1 and PRT/OUR/T1988/1 ASFV strains, and performed haemadsorption assays and immunocytochemistry to monitor infection (Figure 4A, S5). Expression of ASFV proteins p30 and p54, early and late (viral factories), respectively, was detected in the majority of iPSCdMs from all species, indicating that iPSCdMs from the wild and domestic Suidae species supported infection and replication of ASFV equally. Indeed, the levels of virus produced in culture supernatants following infection, as assayed by qPCR for ASFV genomic DNA, were similar to that generated by PAMs and confirmed the active propagation of ASFV in iPSCdMs from all three species (Figure 4B, S5).

**Figure 4.**
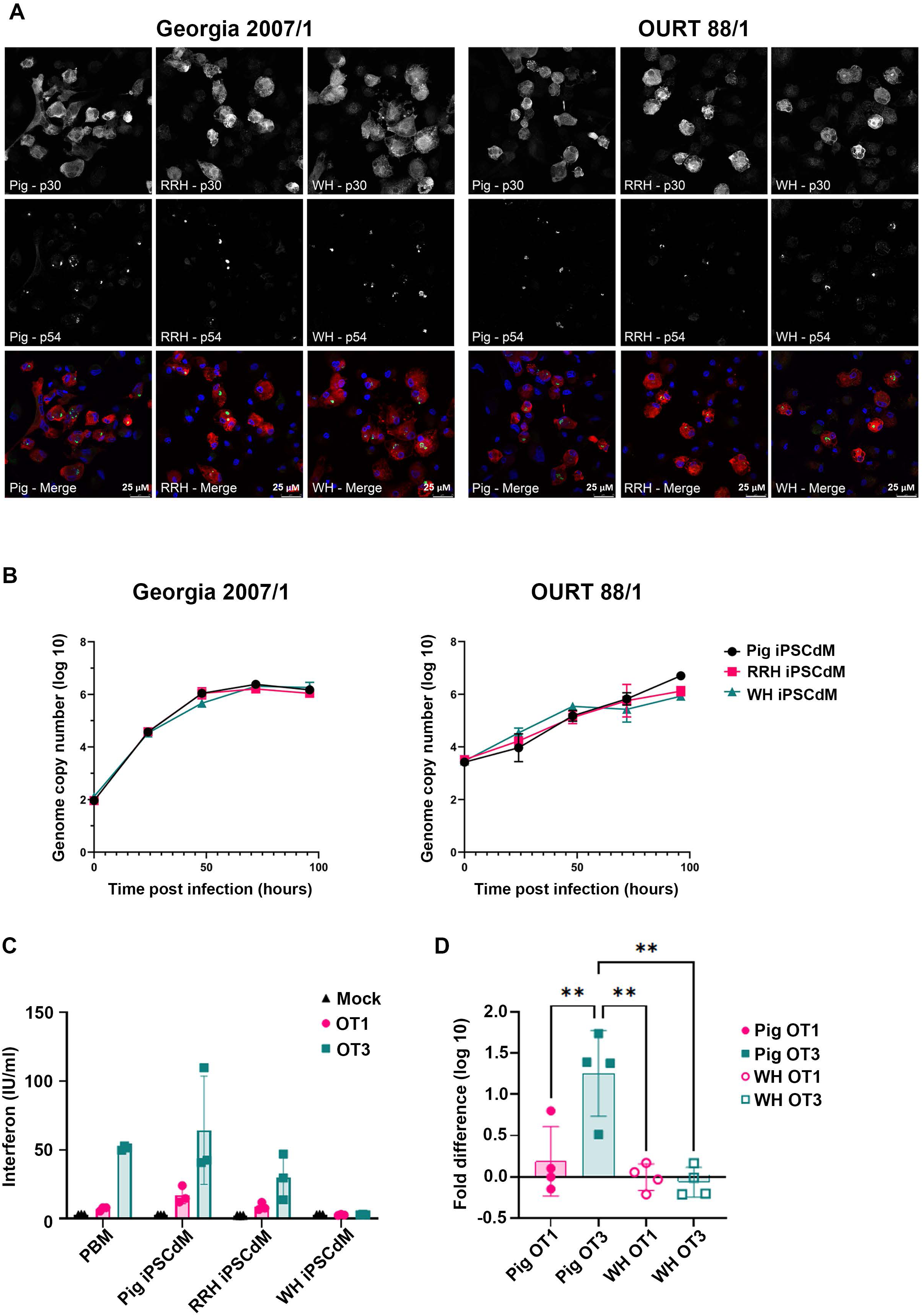
Characterisation of ASFV infection in ASF relevant domestic and wild pig iPSCdMs. (A) Fluorescent image of ASFV-infected iPSCdMs from 3 suid species, 24 hpi, stained for viral proteins p54 (green) and p30 (Red). (B) Pig, red river hog and warthog iPSCdM were infected with Georgia 2007/1 and OURT 88/1 viruses for one hour at a multiplicity of infection of 0.01 and collected at the indicated times. Genome copy numbers were determined by qPCR. Mean and ±SD of 4 technical replicates. (C) Induction of IFN in iPSCdM cells infected with a virulent (OURT 88/1) and attenuated (OURT 88/3) ASFV isolate, measured using SK6 MX2/Luciferase bioassay. Results interpolated using a universal IFNα standard. Mean ±SD of 3 technical repeats. (D) Comparison of porcine and warthog IFN induction expressed as fold differences against mock. Mean ±SD 4 independent experiments and statistical differences determined by one-way ANOVA (** p < 0.01).

IFN production, normally a key early antiviral response, is effectively suppressed in pig Mφs *in vitro* upon infection by ASFV (Dixon et al., 2019; Golding et al., 2016). However, during the later stages of infection *in vivo*, levels of IFN increase to pathological levels, indicating that IFN dynamics might influence development of disease. In line with this, attenuated, less virulent ASFV strains are impaired in their capacity to suppress IFN production (Golding et al., 2016; Reis et al., 2016). To examine how IFN production compared between Suidae species, we infected iPSCdMs with the genotype 1 virulent PRT/OUR/T1988/1 virus and an attenuated PRT/OUR/T1988/3 strain carrying deletions in multigene family (MGF) 360 and 505 that suppress host IFN response (Chapman et al., 2008; Portugal et al., 2020; Reis et al., 2016) (Figure 4C, D). In line with previous reports, IFN production by *ex vivo* Mφs or iPSCdMs was suppressed by PRT/OUR/T1988/1, but stimulated upon infection with PRT/OUR/T1988/3. By contrast, IFN was not produced by warthog iPSCdMs when infected with either the attenuated or virulent ASFV strains. Significantly, a similar pattern of IFN suppression was also obtained with *ex vivo* warthog Mφs indicating that the lack of IFN production was a general feature of warthog Mφs (Figure S5). Sendai virus infection induced IFN expression in all suid iPSCdMs, and confirmed that the warthog Mφs could produce IFN (Figure S5). However, the induction of IFN by OURT88/3 infection in red river hog iPSCdMs may indicate that African wild suid species differ in how they respond to ASFV and manifest resilience.

## Discussion

We previously demonstrated that embryo-derived PSC provide a useful model for studying host-pathogen interactions in the domestic pig (Meek et al., 2022). However, the requirement for embryos to derive new cell lines is normally prohibitive in the case of wild species, and restrictive even for rare domestic breeds. Reprogrammed iPSCs comparable to embryo-derived stem cells therefore represented a useful alternative source of PSCs from different Suidae species. Indeed, the generation of suid iPSC-derived Mφs, displaying molecular and functional characteristics similar to *ex vivo* cells, enables comparative cross-species studies of ASFV resilience. Moreover, suppression of the IFN response in warthog iPSCdMs, in contrast to other suids tested, provides new opportunities to dissect how modulation of this key antiviral response contributes to ASFV resilience in iPSC-derived Mφs.

The transgene-dependent pig iPSCs reported here differentiated readily into Mφs that possessed molecular and functional characteristics equivalent to embryonic PSCdMs. Moreover, iPSCdMs were efficiently infected by ASFV and supported the growth of the virus at levels similar to PSCdMs and *ex vivo* pig Mφs. Although transgene-dependent iPSCs had previously been reported for sheep, pig and cow (Gao et al., 2019; Li et al., 2011; Zhao et al., 2021), providing precedent for this general approach, we were encouraged by the apparent stability of the pig iPSC lines and their broad differentiation potential, despite the fact that chronic high-level expression of the reprogramming factors may not have installed an authentic native pluripotency gene network and epigenetic state. Identification of suitable clones based on their stability and differentiation capacity was effective but necessitated screening significant numbers of clones and did not guarantee efficient and reproducible differentiation. For suboptimal clones, re-expression of the doxycycline-regulated transgenes in iPSCdMs through a second round of reprogramming and re-selection of iPSCs proved useful in improving differentiation potential. Retention of an “epigenetic memory” from the primary iPSCdMs might be responsible for this improvement, as previously hypothesised for resetting mouse iPSC fate (Kim et al., 2010). However, in iPSC clones where reprogramming was heterogeneous and only partially effective leading to differentiation failure or death amongst a sub-population, the second round might simply enrich for differentiation competence per se, rather than biasing differentiation fate.

Inclusion of the histone deacetylase (HDAC) inhibitors sodium butyrate and valproic acid during reprogramming markedly improved the recovery of colonies displaying compact PSC morphology and differentiation potential. This effect on reprogramming might result from increased expression of the doxycycline-induced piggyBac transgenes. However, use of HDACs in a variety of expression systems as well as promoting transition between pluripotent states, supports the notion that the advantages of HDAC inhibitors are mediated through creating a more malleable, programmable, host epigenome (Huangfu et al., 2008; Takashima et al., 2014; Trokovic et al., 2013). HDAC inhibitors were previously reported to improve the generation of pig iPSC clones, but may have compromised differentiation potential (Mao et al., 2017; Petkov et al., 2016). We also noted reduced differentiation efficiency in iPSCs passaged with HDAC inhibitors, but found their potential could be improved following reduction or withdrawal of HDAC inhibitors prior to initiating differentiation. Overall, our results confirm that HDAC inhibition is useful in promoting reprogramming and establishment of Suidae iPSCs, but may require careful tuning to ensure effective differentiation.

The combined use of HDAC inhibitors and piggyBac vector reprogramming enabled the generation of differentiation-competent iPSC lines from different suids. This provided new opportunities to compare ASFV infection and host responses in species displaying different susceptibilities to ASF. Both domestic pigs, wild boar and African wild pig iPSCdMs were readily infected by ASFV and propagated the virus efficiently. Given iPSCdMs represent an embryonic cell type, albeit one that contributes to tissue resident Mφs in the adult, iPSCdMs might be assumed to have a relatively naïve phenotype endowed by their derivation in culture. Nevertheless, despite their distinct origins, the similar efficiencies of ASFV propagation in pig iPSCdMs and *ex vivo* Mφs demonstrated that iPSC-derived cells should provide a useful cell culture model for investigating ASFV-host interactions. This conclusion was further supported by the comparable patterns of IFN production when infected by virulent and attenuated ASFV strains.

In contrast to the other Suidae iPSCdMs tested, warthog iPSCdM and *ex vivo* Mφs did not induce significant levels of IFN when infected with the attenuated ASFV. This could reflect an evolved adaptation in warthogs to moderate the antiviral response, facilitating immune tolerance and resilience to ASFV. Dampening the inflammatory response may avoid triggering the cascade that leads to chronic cytokine release and ensuing pathological damage in domestic pigs. Bats, the archetypal models of virus tolerance, are reservoirs for many different viruses and have been reported to adopt a number of strategies to limit an inflammatory response (Irving et al., 2021). For example, Nipah and Hendra virus are endemic in the Large flying fox (*Pteropus vampyrus*), but do not induce an IFN response, nor cause disease (Glennon et al., 2015). The distinct response in warthog may indicate that despite the deletion of key genes that enable IFN induction in the other suids, ASFV may possess additional factors that suppress IFN activation in warthogs, or warthog Mφs may exhibit a particular response to the pathogen-associated molecular patterns produced by ASFV.

By contrast, the increased IFN production in red river hog iPSCdMs infected with attenuated ASFV indicates that suppression of IFN is not necessarily an essential feature associated with ASFV resilience. Indeed, reduced growth of ASFV was reported in SV40 large T immortalised red river hog Mφs, indicating that limiting virus replication could be a contributory factor for resilience in this African suid (Takenouchi et al., 2024). The immortalised red river hog Mφs were generated from blood monocytes expanded in the presence of kidney cells and therefore may reflect a particular subtype of Mφs or a particular immune/polarisation state. In addition, phenotypes of blood monocytes have only been defined for the pig in any detail (Chamorro et al., 2005), and it is possible that different populations of monocytes may circulate in the blood of other suids. We occasionally observed reduced infectivity of red river hog iPSCdMs in less efficient and more heterogeneous iPSC differentiations. This variability was not observed with pig iPSCdMs and could indicate that red river hog iPSCdMs are more readily triggered to adopt an ASFV-resistant state.

In summary, we have generated a set of suid iPSC lines that display different responses to ASFV that may have relevance to understanding natural resilience. The derivation of iPSCs from endangered Suidae species, such as the Visayan warty pig (*Sus cebifrons*) and members of the Babirusa (*Babyrousa*) genus, could also provide insights into the basis of their vulnerability to ASFV. Whilst iPSCdMs are not strictly equivalent to *ex vivo* Mφs, their use should enable investigation of how host genetics controls ASFV resilience and resistance through gene editing in iPSCs and iPSCdMs. Moreover, PSCs can differentiate into other cell types that play a role in development of ASF disease, such as dendritic, natural killer, and endothelial cells. In this way, Suidae iPSCs can enable engineering multi-cellular, even cross species, co-culture systems to interrogate the complex cell-cell interactions that operate during progression of ASFV infection and disease.

## Methods

### Sources of primary cells for reprogramming

Primary pig embryonic fibroblasts were derived from two Large White (CMV-EGFP) / Landrace (wild-type) male embryos at ∼day 35 of embryonic development (A. J. Thomson et al., 2012). Male wild boar fibroblast-like cells were established from a small population of adherent, fibroblast cells present in a preparation of wild boar pulmonary alveolar Mφs (PAMs). The PAMs were sourced from a slaughterhouse following the processing of a Scottish farm-reared adult wild boar. Red river hog fibroblasts were derived from discard scrotal skin tissue derived at the Royal Zoological Society Scotland Edinburgh Zoo through routine castration of two young male siblings (ZBQ16-06210, ZBQ16-06211). Both animals were born at the Royal Zoological Society Scotland’s Edinburgh Zoo and samples were supplied to the Roslin Institute under Material Transfer Agreement RTA2475. Two male warthog fibroblast cultures derived from either trachea or brain provided by the Collection of Cell Lines in Veterinary Medicine/Department of Experimental Animal Facilities and Biorisk Management at the Friedrich Loeffler Institute under Material Transfer Agreement ZB22-214.

### Reprogramming and passaging of suid iPSCs

Suid fibroblasts were resuspended in Resuspension Buffer R (Invitrogen) at 0.5-1×10^6^ /100 µl and electroporated using the Neon^TM^ Electroporation System (Invitrogen) set at: 1650V, 10ms pulse width and 3 pulses. Electroporated cells were seeded at 2.4×10^3^ cells/cm^2^ on γ-irradiated mitotically inactive STO feeders (4×10^4^/cm^2^) in 6-well tissue culture plates in M15G+NaB medium [G-MEM (Sigma, #G5154), 15 % FBS (Gibco, #10270-106) 1x NEAA (Gibco, #11140035), 1 mM sodium pyruvate (Gibco, #11360039), 2 mM L-glutamine (Gibco, #25030024), 0.1 nM β-mercaptoethanol (Gibco, #31350010), 1000 U/mL hLif (R&D, #7734-LF), 50 µg/mL L-ascorbic acid (Sigma, #49752-10G), 1 µg/mL doxycycline (Sigma, #D9891-5G), 0.5-1 mM sodium butyrate (Aldrich, #303410-5G) and cultured at 37 °C in a 5% CO_2_ incubator. Warthog M15G reprogramming media contained 0.5 mM sodium butyrate and 0.3 mM of valproic acid (Sigma #P4543-10G). The media was changed every other day and iPSC-like colonies emerged at 10-14 days. At 13-30 days suitable iPSC colonies were manually dispersed by pipetting, transferred to 96-well plates containing STO feeders and subsequently expanded by passaging. Routinely, iPSC cultures were dissociated using 0.025% trypsin/EDTA (Trypsin: Life technologies, #25200056) in PBS containing 1mM EDTA) and sub-cultured at 2-3 × 10^3^ / cm^2^ in M15G+NaB medium with 0.5 mM sodium butyrate and refed daily. Typically, iPSCs were passaged every 3–5 days. Experiments typically used iPSC lines at <20 passages, and cultures were regularly tested to confirm they were mycoplasma free.

### Karyotype analysis

Cells with 0.1 mg/ml KaryoMAXTM ColcemidTM (Thermo Fisher: #15212012) solution in M15G (37 oC, 5% CO2) for 2-3 h, swollen in 0.56% (w/v) KCl hypotonic solution for 10-15 minutes at room temperature and fixed with 1/50 volume of −20^0^C 3:1, methanol:acetic acid fixative. Cell pellets were resuspended in fixative, incubated on ice for 30 minutes, pelleted again and resuspended in fixative prior to dropping onto glass slides to generate Metaphase spreads. Slides were dried overnight, stained with KaryoMAX Giemsa Stain (Thermo Fisher: #10092013) and DAPI (Abcam: ab228549). 20 metaphase spreads were assessed per cell line by light microscopy and chromosomes were counted using ImageJ. Chromosome counts were Pig iPSC1 14/20 diploid (38); RRH iPSC1 15/20 diploid (34); and Warthog iPSC 15/20 diploid (34).

### Alkaline phosphatase staining

Alkaline phosphatase activity was detected in situ using the Alkaline Phosphatase Detection Kit (Sigma #SCR004) according to the manufacturer’s instructions.

### Immunostaining

Cells were fixed in 4 % Paraformaldehyde at room temperature (RT) for 10 min, washed 3 x 5 min in PBS, and permeabilised with 0.5 % v/v TritonX100/PBS for 30 min. Fixed samples were blocked with 1 % BSA (Life Tech, #15260037), 1 % donkey serum (Merck, #D9663) and 1 % goat serum (Merck, #G6767) for 30 min and then incubated overnight at 8 °C, with either EPCAM PE (1/100) (Thermo Fisher, #12-5791-81), OCT4 (1/100) (Santa Cruz, #C-10:sc-5279) or NANOG (1/100) (Abcam, #ab80892). Samples were washed 3 x with 0.1 % Tween 20/PBS and incubated with secondary antibody (OCT4: Alexa Fluor 546 goat α-mouse (1/500), NANOG: Alexa fluor 488 goat α-rabbit (1/500)) for 1 h at RT. Samples were washed 3 x 0.1% Tween 20/PBS, with final wash containing (1/100) DAPI, and imaged. Neural differentiation was confirmed by beta3-Tubulin staining. Samples were permeabilised by addition of ice-cold methanol and incubated at −20°C for 10 min, washed 3 x with PBS for 1-2 min and blocked for 1 h at RT using PBS-T (0.1% v/v TritonX100) containing 10% goat serum (Merck, #G6767). The blocking solution was replaced with primary TUJ1 antibody (Covance, #mmms-435P), diluted 1/500 in blocking solution. The plates were incubated overnight at 4°C and washed 4 x with PBS-T for 5 min. The secondary antibody (Goat α-mouse AF546, 1:1000) was diluted in blocking solution and incubated on samples for 1 h at RT in the dark. Plates were washed 4 x with PBS-T, with DAPI (1:10,000 in PBS-T), and 2 x PBS further washes before imaging.

### RT-qPCR

RNA was prepared using Qiagen RNeasy kit (#74104) following the manufacturer’s protocol including on-column DNase treatment. cDNA was synthesised from 0.2–1 μg of RNA using Agilent’s Multitemp cDNA Synthesis kit (#200436) at 42°C following the manufacturer’s instructions. cDNA samples were diluted to 1 μg / mL with nuclease-free water. Each RT-qPCR reaction consisted of 8 μl of diluted cDNA plus a master mix consisting of 10 μl Agilent Brilliant III SYBR green (#600883), 0.4 μl Reference dye (2 μM) and 0.8 μl each of forward and reverse primers. *RPL4* housekeeping gene primers were used to normalise expression. The reaction was performed using a Stratagene MxPro3005P set at: one cycle of 95°C for 2 min, 40 cycles of 95°C for 15 s and 60°C for 30 s. A final cycle of 95°C for 1 min, 60°C for 30 s and 95°C for 15 s was used to establish a dissociation curve.

### Directed iPSC differentiation

For neuronal differentiation 4000 iPSCs/well were aliquoted into V-bottom 96 wells in 200 μl of N2B27 media [DMEM/F12 (Gibco, #21331-020)/ Neurobasal (Gibco, #21103-049) (1:1) supplemented with 1x N2 (Thermo Fisher, #17502048), 1x B27 (Thermo Fisher, #17504044), 2 mM L-Glutamine, 0.1 mM β-mercaptoethanol, 20 ng/mL], with EGF (Peprotech,# AF-100-15), 20 ng/mL and bFGF (Qkine, #Qk027), and aggregated by centrifugation at 1000 rcf for 3 min. EBs were fed 2 days post aggregation and after 4 days plated onto laminin (Merck, #L2020) precoated (10 μg/mL for 3 x h at 37°C) culture dishes in N2B27 media. Medium was replaced every 4 days until being fixed (4 % Formaldehyde for 15 min) on day 28.

For adipose differentiation iPSCs were plated on 1 µg/mL bovine fibronectin (Merck #F1141) coated tissue culture plastic at 4×10^4^/ cm^2^ in basal bone and chondrocyte (BBC) media [High glucose DMEM (Life Tech #41966), 10% FCS, 1x NEAA, 1x Glutamax (Life Tech #35050061), 1x Pen/Strep (Life Tech #15140122), 0.1 nM β-mercaptoethanol, 60 µg/mL L-Ascorbic acid, ITS-X (10 µg/mL Insulin, 5.5 µg/mL Transferrin, 0.0067 µg/mL sodium selenite and 2 µg/mL Ethanolamine (Life Tech #51500056)]. At 4 days post plating the media was changed to Adipose differentiation media (BBC media with addition of 500 µM IBMX (Sigma #I5879), 1 mM Dexamethasone (Sigma #D4902), 1mM Rosiglitazone (A XON MEDCHEM #2443) and 0.2 nM 3’ 3’ 5’ triiodo-l-thyronine (T3) (Tocris #6666). Media was replaced every 2-3 days, and on day 21 cells were stained with BODIPY (Invitrogen, #D3922) (Qiu & Simon, 2016)and imaged.

### Stable transfection and gene editing of iPSCs

Stable transfection and *REX1* gene-targeting was performed using CAG-EGFP-IRES-puro and *REX1*-EGFP targeting constructs, respectively, as described previously (Meek et al., 2022). The CRISPR/Cas9 guide sequence (Rex1_363 CTTCTTTCACTGATTTGTAT (Synthego) is conserved between domestic pigs and red river hog and directs cleavage 8 bp upstream of the *REX1* stop codon. Ribonucleoprotein complexes (RNPs) were formed by combining 7.5 µl sgRNA (100 µM) and 1.6 µl Cas9 protein (62 µM, IDT, #1081059) for 10 min at RT. Prior to transfection, 1 µg of the targeting vector (Meek et al., 2022) was combined with the RNPs and diluted in 100 µl of Resuspension Buffer R on ice. 5 ×10^5^ iPSCs were resuspended in the Resuspension Buffer R / RNP / targeting vector mix and electroporated with the Neon^TM^ transfection system using parameters: 1100 V, 30 ms pulse width, 1x pulse. Electroporated iPSCs were resuspended in M15G+NaB and plated in 2 wells of a 6-well plate on mitotically inactive STO feeder cells. The medium was replaced with M15G+NaB the next day, and 72 h after transfection iPSCs were passaged and plated onto feeder cells at 1×10⁴ cells/cm². 24 h after plating medium was replaced with M15G+NaB medium containing 1 µg/mL puromycin and replaced daily until puromycin-resistant colonies emerged (≥ 9 days). Domestic pig and red river hog clones were picked and expanded, and screened by genomic PCR for correct insertion of the targeting cassette at both the 5’ and 3’ sides of the integration site using the following conditions: 1. 98°C for 1 min; 2. 32 x 98°C for 30 s, 69°C for 30 s, 72°C for 1 min; 3. 72°C for 10 mins. PCR products of ∼885 bp and ∼1213 bp identified correct insertion at 5’ and 3’ends respectively.

### Mφ differentiation

PSC/iPSCs were differentiated into Mφs essentially as described previously (Meek et al., 2022). Typically, 3000–6000 iPSCs were dispensed per well into a 96-well V-bottomed plate containing 100 μl mesoderm induction medium [StemPro (Thermo, #A1000701), 25-50 ng/mL rhBMP4 (R&D, #314-BP), 50 ng/mL rhVEGF (R&D, #293-VE), 20 ng/mL and rhSCF (R&D, #255-SC)] and centrifuged at 1000 rcf for 3 min. The aggregated iPSC EBs were refed with 100 μl fresh mesoderm induction medium on days 1 and 3 after aggregation. On day 4, 10–20 EBs were transferred to a gelatinised 6-well tissue culture plate containing Mφ Induction media (X-Vivo 15 (Lonza, #LZBE02-060F), 2 mM Glutamax, 50 nM β-mercaptoethanol, pen/strep, 100 ng/mL recombinant porcine M-CSF (Roslin Technologies), and 25 ng/mL rpIL-3 (Kingfisher Biotech, #RP1298S)]. Attached EBs were fed every 4 days with Mφ induction medium. Floating clusters of round cells with small projections were normally observed in the medium after 9-12 days, signifying early signs of Mφ differentiation. Typically, these relatively immature Mφs were first harvested from culture medium at about day 20 and collected every 4 days thereafter until day ∼48. Mφs were matured by plating cells on non-coated tissue culture dishes in Mφ maturation medium (X-Vivo 15, 2 mM Glutamax, pen/strep, 100 ng/mL rpM-CSF).

### Phagocytosis assay

iPSCdMs were plated at 5-10×10^4^ cells/cm^2^ in Mφ maturation medium for 48 h prior to the addition of 100 μg/mL sonicated (70 kHz for 30s to dissociate beads) pHrodo Red Bioparticles (Thermo, #P35364). The iPSCdMs were imaged 3 h after pHrodo addition.

### LPS induction

iPSCdMs were plated at a density of 5×10^4^/cm^2^ in Mφ maturation medium for 48 h prior to induction with LPS (200 ng/mL Lipopolysaccharides from *Escherichia coli* O111:B4, Sigma #L4391) and harvested after 6 h for RNA preparation.

### Secondary iPSC reprogramming

Mature iPSCdMs were cultured in Mφ maturation medium for 24 h on bacterial culture plates to remove any non-adherent contaminating cells and then replated onto mitotically inactive STO feeder layer in M15G+NaB (containing doxycycline) at 0.2-1×10^4^ cells/cm^2^. The medium was replaced every two days and colonies with iPSC-like compact morphology emerged within 7-14 days. These secondary iPSC colonies were picked, broken into smaller clumps, and plated into wells of a 96-well plate containing feeder cells, and then expanded by passaging as described previously.

### Cell surface staining of Mφs and flow cytometry

Cells were stained with antibodies as described previously (Meek et al., 2022). Cells were blocked with 2% FBS/PBS. Antibodies were diluted in the block solution and incubated with cells for 30 min on ice, in the dark; and resuspended 1:1 PBS/SYTOX Blue Nucleic Acid Stain (5 μM, ThermoFisher #S11348) immediately prior to flow cytometer analysis to allow live/dead cell discrimination. Antibodies used were CD14 (Biorad, #MCA1218F, 1:50) with isotype control (Sigma, #SAB4700700), CD16 (Biorad, #MCA1971PE, 1:200) with isotype control (Biorad, #MCA928PE), CD163 (Biorad, #MCA2311F, 1:100) with isotype control (Sigma, #F6397), CD169 (Biorad, #MCA2316F, 1:100) with isotype control (Sigma, #F6397) and CD172a (Southern Biotech, #4525-09, 1:400) with isotype control (Biorad, #MCA928PE). Flow cytometry analysis was performed using a BD LSR Fortessa (16 colour Analyser) with a High Throughput Sampler.

### PRRSV infection and staining

iPSCdMs were infected with PRRSV strain SU1-Bel (MOI: 1) in cRPMI+CSF1 for 2 h at 37 °C / 5% CO2 as described previously (Meek et al., 2022). After virus removal and recovery overnight in fresh cRPMI+CSF1, iPSCdMs were harvested with a cell scraper, fixed in 4% Formaldehyde/PBS, permeabilised with 0.1% TritonX100/PBS, and incubated with the primary antibody (Rural Technologies #SDOW-17A, 1:5000 in blocking solution) followed by the fluorescent secondary antibody (Goat α-mouse AF488, 1:5000). PRRSV infection was measured by flow cytometry.

### ASFV Infection, staining and growth assays

The ASFV infections performed at APHA (Figure 2H) were performed as described previously (Meek et al., 2022), using ASFV strain Armenia 07 at MOI of 1. Viral DNA levels in post-infection culture supernatants were quantified by RT-qPCR analysis of ASFV VP72 gene expression. ASFV viral genome copy numbers were determined by comparing the Cq expression values to a defined standard produced by multiple dilutions of the pASFV-VP72 plasmid.

The remaining ASFV infections were performed at The Pirbright Institute. Virus cultures with African swine fever virus were carried out in the high containment laboratories at the Pirbright Institute in the United Kingdom of Great Britain and Northern Ireland, under a licence granted by the Health and Safety Executive under Article 4(1) of The Specified Animal Pathogens Order (SAPO) 2008 (statutory instrument 2008/944). All Pirbright Institute ASFV virus stocks were grown in primary porcine bone marrow cells and mock inoculum were prepared in parallel from uninfected cells. GEO2007/1, PRT/OUR/T1988/1, PRT/OUR/T1988/3 and mock were precleared using Vivaspin columns (1,000,000 kDa molecular weight cut-off; Sartorius Stedim Biotech) to remove endogenous cytokines. Sendai virus was purchased from Charles River Laboratory.

To assess viral replication iPSCdMs were infected with either Georgia 2007/1 or PRT/OUR/T1988/1 at an MOI of 0.01 in triplicate. DNA was extracted from supernatants and cells after freeze/thawing twice at different times post infection using MagMax CORE nucleic acid purification kit (ThermoFisher # A32702). qPCR was carried out on an Agilent AriaMx system using Brilliant III UF MM QPCR/ Low ROX High performance (Agilent Technologies, #600890) following a protocol modified from that described by King and coworkers (King et al., 2003) using the primers VP72 sense (CTG CTC ATG GTA TCA ATC TTA TCG A), VP72 antisense [GAT ACC ACA AGA TC(AG) GCC GT] and the probe 5′-(6-carboxyfluorescein [FAM])-CCA CGG GAG GAA TAC CAA CCC AGT G-3′-(6-carboxytetramethylrhodamine [TAMRA]). Quantification of genome copy numbers were derived from a VP72 plasmid standard curve.

For imaging iPSCdM cell monolayers were grown on poly-D-lysine (Thermofisher, # A3890401) coated glass coverslips and incubated for 3 days. The cells were infected with a high MOI for 1 hour and the inoculum removed and replaced with fresh media. At 18 hpi the cells were fixed with 4% paraformaldehyde (Merck, #P6148) for 1 hour. The cells were permeabilised using 0.2% Triton-X-100/PBS for 5 min and blocked for 30 min in PBS/10% normal goat serum (Merck, #S26)/0.2% fish skin gelatin (Merck, #G7041). Primary antibodies, anti p72 and anti p30 were diluted in block, added to the cells, incubated for 1 hour, and washed 3 times with PBS/0.2% tween 20. Secondary antibodies, anti-rabbit Alexa Fluor 488 and anti-mouse Alexa Fluor 568 (Thermofisher, #A-11008 and #A-11004 respectively) were added diluted 1/500 in blocking buffer and incubated for 1 hour. After washing 3 times, DAPI (4′,6-diamidino-2-phenylindole: Merck, #MBD0015) diluted 1/10,000 in water, was added for 5 min. The coverslips were mounted on glass slides using Glass Antifade Mountant (Thermofisher #P36982) and visualised using a Leica SP8 laser scanning confocal microscope.

For Hemadsorption assays iPSCdMs were infected with a high MOI of virus and incubated overnight at 37^0^C. Porcine blood cells were washed twice with PBS and added to the infected cells and incubated for a further 16 hours. Formation of rosettes was observed and imaged using a Leica DMi1 microscope attached to a Leica MC170 HD camera.

### IFN Induction Assay

The *Mx2* promoter was amplified from pig genomic DNA using primers 5’-NNNNNNACGCGTGACCTAACTGGCTGGATCT and NNNNNNCTCGAGCCGAGCTCTTACCTGAGCAC and cloned into the *Mlu* I and *Xho* I sites of pGL3-Basic. An *Mx2* promoter-firefly luciferase cassette was then subcloned into pcDNA3 in place of the CMV promoter using *Mlu* I and *Bam*H I restriction enzymes. All inserts were confirmed by Sanger sequencing. A stable SK6 cell line expressing firefly luciferase under the control of a porcine *Mx2* promoter was then cloned and expanded in the presence of 500 µg/ml G418. Samples, along with a two-fold serial dilution of recombinant universal IFNα1 (PBL assay Science, #11200) starting at 500 IU/ml down to 3.9 IU/ml were incubated on SK6-Mx2-luc cells to quantify secreted IFN.

iPSCdM cells were plated on 48 well plates in triplicate for each treatment and incubated for 3 days. The cells were infected with pre-cleaned viruses at an MOI of 2 in triplicate and the mock included. 25 U per well Sendia virus (Charles River laboratory) was included as a positive control for IFN induction. After 24 hours the supernatants/IFN were transferred to plates with SK6-Mx2-Luc cells. After 24 hours at 37^0^C the supernatant was removed and cells were lysed using Glo-lysis buffer (Promega #E2661) and transferred to white 96 well plates. An equal volume of Bright-Glo reagent (Promega, #E2610) was added and incubated for 5 min. The plates were read on a Promega GloMax Discovery microplate reader, using Bright-glo program.

### RNA Sequencing Analysis

Total RNA was prepared for three technical replicates of domestic pig 2 iPSC and iPSCdMs and starting PEFs, four technical replicates of RRH iPSCs and iPSCdMs, two technical replicates of warthog iPSCdMs and 1 sample of wild boar iPSC, iPSCdMs and warthog BMDMs. RNA samples were extracted using the Qiagen RNeasy Kit (#74104) in accordance with the manufacturer’s instructions, including the recommended on-column DNase treatment. Short-read RNA-seq libraries were generated with the TruSeq Stranded mRNA Library Prep Kit (Illumina). Briefly, poly(A)+ mRNA was isolated and fragmented, after which first-strand cDNA synthesis was performed using reverse transcriptase and random primers. Libraries were sequenced on an Illumina NovaSeq platform to produce 2×100 bp paired-end reads.

For the comparative transcriptomics analysis, a total of 24 fibroblast, iPSC and iPSC derived macrophage (iPSCdM) samples from Suidae species (pig, wild boar, red river hog and warthog) were sequenced using Illumina RNA seq and deposited in ENA under accession PRJEB111747. Additionally, 31 RNA seq samples from diverse tissues were downloaded, including three macrophage samples from a Duroc pig (PRJEB19386;(Warr et al., 2020)). Pig glia expression was calculated from 15 samples (PRJNA722748; (Shih et al., 2023)). Stem cell and stem cell derived macrophage data were obtained from PRJNA787759 (Meek et al., 2022).

### Bioinformatics

RNA-seq data analysed included those previously reported (Meek et al., 2022) and additional wild and domestic pig iPSC and iPSCdMs, domestic PEFs and warthog BMDM datasets. Publicly available pig tissue and cell line specific RNA-seq datasets were downloaded from ENA (microglia: PRJNA722748, ESC and ESCdMac: PRJNA787759 and pig somatic tissues and cell lines: PRJEB19386). The paired-end Illumina short-read data were adapter trimmed with cutadapt (v.2.10) with parameters “-m 30 -O 10 –match-read-wildcards“ using the standard Illumina adapter sequences. Trimmed reads were mapped to the pig reference genome (Sscrofa11.1) using STAR (v 2.7.1a) only allowing a maximum of 10 multimappers per read. Mapping rates were consistently above 90%, apart from the iPSC data, for which it dropped below 70%. This is likely due to the mouse feeder cells which are required to maintain the stem cell cultures. Gene level read counts were generated with featureCount (v. 1.6.3) [60] using the Ensembl pig genome annotation (v.101) [61]. Heat map, sample specific clustering and PCA plots were created in R (https://www.R-project.org/.) using the DESeq2 package [60]. Genes of low expression were filtered out (total read counts per gene < 20), and a variance stabilising transformation was used before comparing the samples.

### Species identification

SNPs in suid *RELA* genes distinguish domestic pigs, warthogs and red river hogs (Palgrave et al., 2011) (Figure S6). RELA genomic DNA fragments were generated from the suid iPSCs by PCR with the following cycle conditions 1. 98°C for 1 min. 2. 32 x (98°C for 30 s, 65°C for 30 s, 72°C for 30 s). 3. 72°C for 2 min. For sequencing 500-750 ng of purified DNA and 6.4 pmol of sequencing primers (For: GGAAGGGACACTGACAGAGG, Rev: TCAGAAGGGCTGAGAAGTCC) were combined and made up to a total volume of 6 μl with nuclease free ddH_2_O. Sequencing was performed by Edinburgh Genomics using a BigDye v3.1 Terminator Cycle Sequencing Kit (Thermofisher P/NAB0384/240; BS034042) and analysed on a 3730xl DNA Analyzer (Applied Biosystems™, 3730xl).

Additionally, RT-qPCR using established primers (RPL4 Species ID and NANOG UTR), designed for domestic pig, can distinguish between the three species due to differences in melt curve temperatures (Figure S6).

### Ethical approval

Bone marrow cells were obtained from pigs humanely killed at The Pirbright Institute under the auspices of Establishment License X24684464. The process was controlled by the Animal Welfare and Ethical Review Board (AWERB) of The Pirbright Institute. The collection of tissue samples and derivation of cell lines was approved by the Veterinary Ethical Review Committee at the Royal (Dick) School of Veterinary Studies, University of Edinburgh.

### Statistical analysis

Data was analysed using Prism v10.4.0 (GraphPad Software, LLC). Fold-differences in interferon induction were compared using 2-way ANOVA on log transformed data after normality was assessed using the Shapiro-Wilk test.

### Resource availability Corresponding author

Requests for further information, resources, and reagents should be directed to the lead contact, Tom Burdon

## Materials availability

All iPSC lines generated in this study are available and will be supplied under a materials transfer agreement upon request from the corresponding authors, Royal Zoological Society of Scotland or Fredrich Loeffler Institute as necessary.

## Data and code availability

All data generated in this study adhere to FAIR (Findable, Accessible, Interoperable, and Reusable) principles, and data sharing respects CARE (Collective Benefit, Authority to Control, Responsibility, and Ethics) principles. All raw data and associated processed datasets, including characterization data for all iPSC lines, are available from the lead contact upon reasonable request.

## Supporting information

Watson et al Supplementary figures

## Acknowledgments

The authors acknowledge the support of Roslin Technology Limited for Tom Watson during his PhD study and BBSRC Impact Accelerator Award to the University of Edinburgh PIII054. The authors also thank Dr Ian Valentine and Dr Christine Tait-Burkard for providing tissue/cell samples and PRRSV virus stock. This work was supported by funding from the Biotechnology and Biological Sciences Research Council Institute Strategic Programme Grants BBS/E/D/10002070; BBS/E/D/10002071; BBS/E/RL/230001C; BBS/E/RL/230002A; BBSRC EASTBIO CASE PhD studentship; and National Centre for the Replacement, Refinement and Reduction of Animals in Research (NC3Rs) grant NC/V001140/. LCG and CLN are supported by BBSRC grants BBS/E/PI/0000230002B and BBS/E/PI/000023NB0004, and would like to acknowledge the Pirbright Institute’s Bioimaging Science Technology Platform that is also supported through UKRI grant BBS/E/PI/23NB0004.

## Author contributions

TMW, LCG, SM, SK, and NB performed the experiments. S.S, MJ, AF and IH generated karyotype data, Immunocytochemistry and images. ALA contributed resources for RNA sequencing and LE performed the bioinformatic analysis. SK and SG provided the tissue samples. TB, TW, CLN, and JM conceived the study. FS, HC, CLN, FG and TB directed the study. TB and TW wrote the manuscript with input from CLN, LCG, SK, SG, ALA, FG and FS.

## Supplemental information

Document S1. Figures S1-S6 and Table S1.

