## Supplementary material for "Suidae iPSC-derived macrophages as models for investigating susceptibility and resilience to African swine fever virus": Watson et al Supplementary figures

**Document S1**

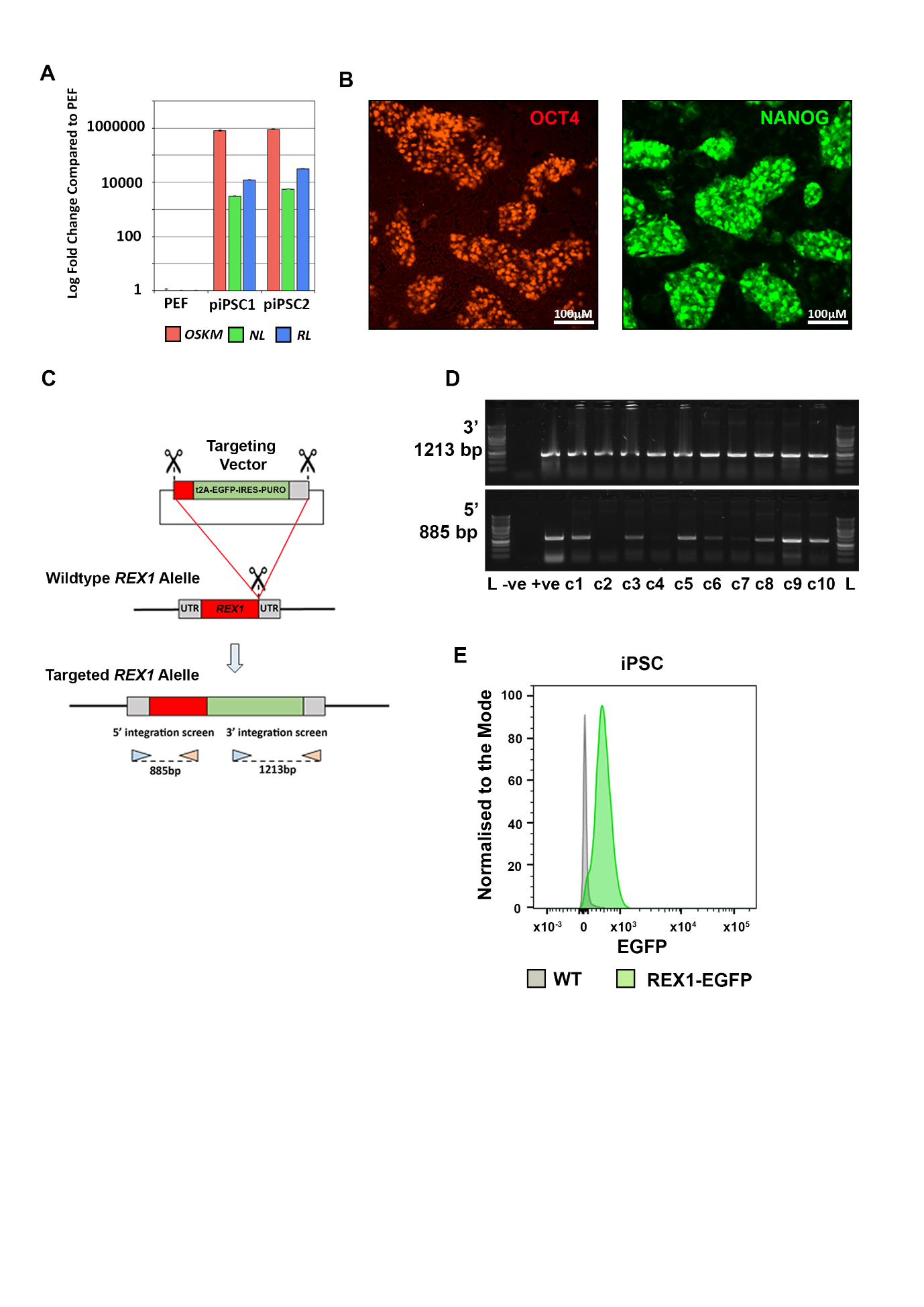

**Figure S1** **Characterisation of domestic pig iPSC pluripotency factor expression.** (A) RT-qPCR expression data showing the log fold change of reprogramming plasmid derived mRNA in piPSC1 and piPSC2 relative to non-transfected fibroblasts. Mean ±SD of three independent experiments. (B) Fluorescent images of OCT4 (Red) and NANOG (Green) protein expression in piPSC1. (C) Diagram showing *REX1-EGFP* Knock-in strategy. CRISPR/Cas9-mediated homology-directed repair, using a repair template (top), allows the generation of the targeted *REX1* allele. PCR product sizes for integration are indicated. (D) PCR genotyping of puro-resistant, EGFP+ clones. Clones 1, 3, 5, 6, 8, 9 and 10 showed the expected products at both the 5’ and 3’ ends of the integration site. Wild-type, parental, porcine iPSC genomic DNA was used as a negative control (-ve) and the transfected pool as a positive control (+ve). (E) Flow cytometry analysis of porcine *REX1-EGFP* iPSCs and iPSCdMs.

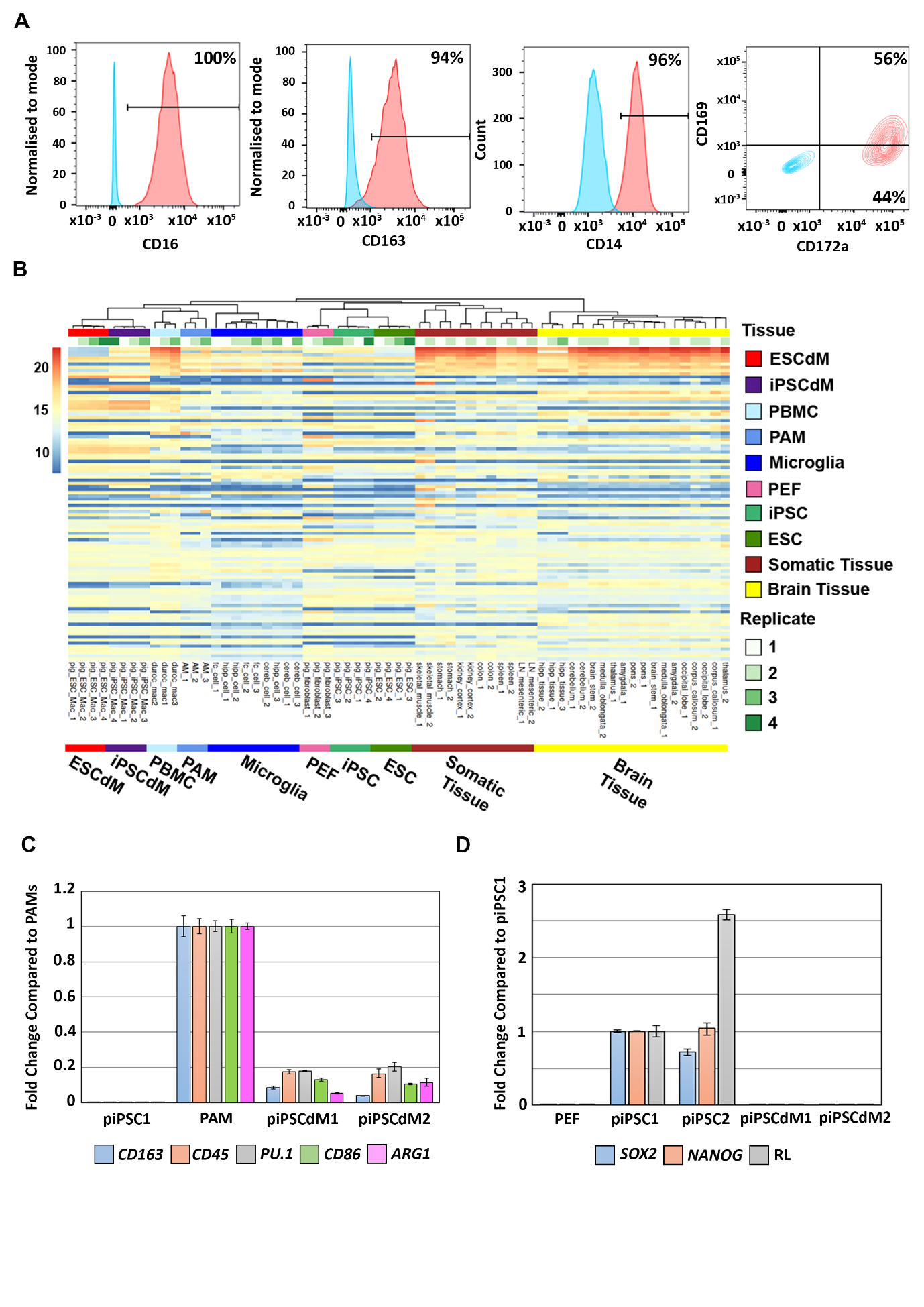

**Figure S2** **Additional characterisation of piPSC1, piPSC2 and derived iPSCdMs.** (A) Flow cytometry of piPSC2 derived Mφs stained for Mφ surface markers (Red) relative to isotype controls (Blue). (B) Heatmap showing the hundred most highly expressed genes in cell lines and pig tissues. The tree at the top of the heatmap indicates the hierarchical clustering between samples. (C) RT-qPCR data showing expression of endogenous Mφ genes in iPSCdMs compared to undifferentiated piPSCs and *ex-vivo* PAMs. Mean ±SD of 3 independent experiments. (D) RT-qPCR data showing the total (endogenous + exogenous) expression of pluripotency genes *SOX2* and *NANOG*, and the *RARG+LRH1* reprogramming transgene in piPSCs compared to fibroblasts and iPSCdMs. Mean ±SD of 3 independent experiments.

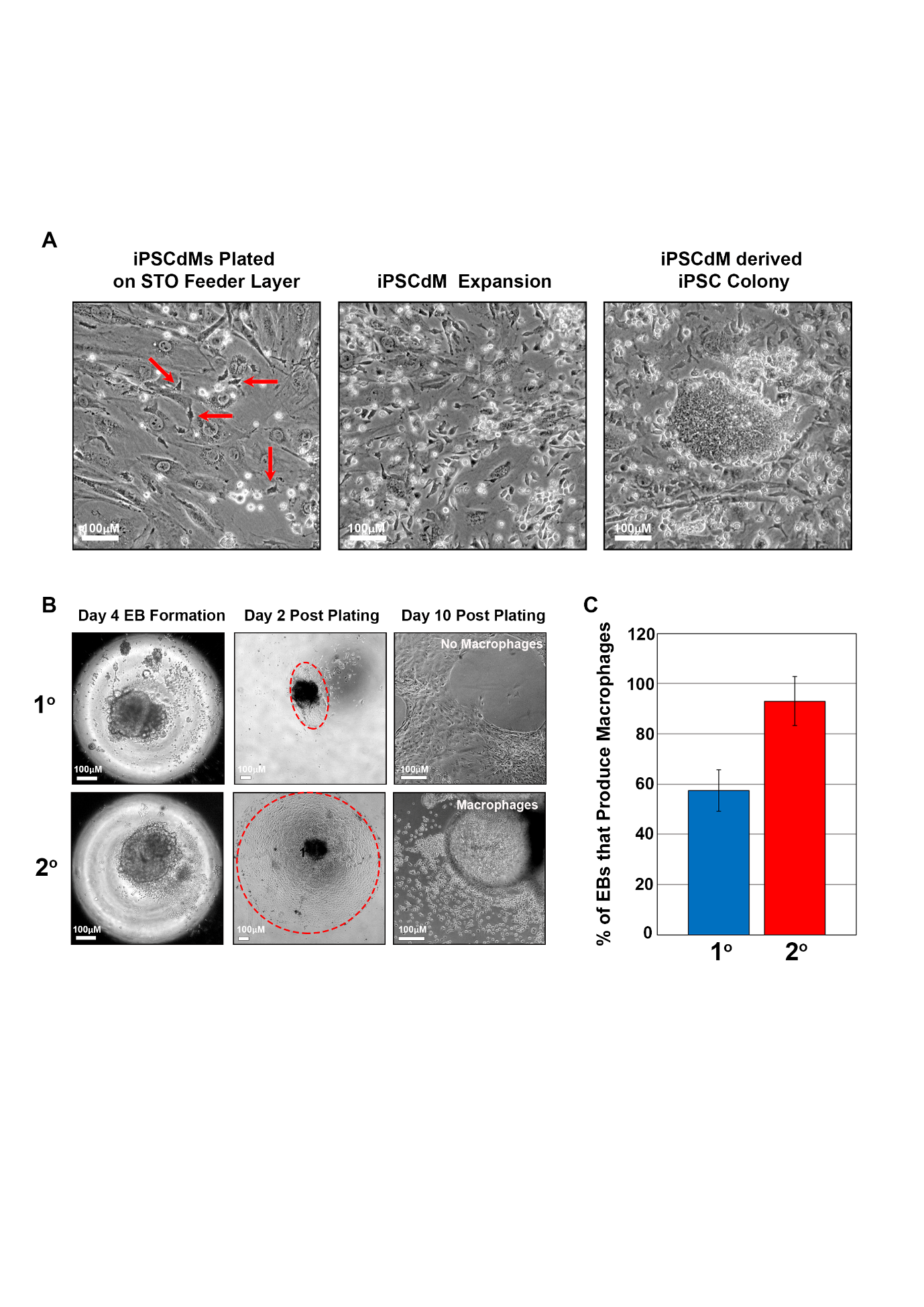

**Figure S3** **Secondary reprogramming of suid iPSCs to improve Mφ differentiation potential.** (A) Representative bright field images showing the stages of iPSCdM secondary reprogramming. Red arrow indicates iPSCdMs adhered to feeder layer. (B) Representative bright field images showing the effect of secondary reprogramming on EB outgrowth and Mφ production. (C) Bar graph showing the mean percentage of EBs capable of producing Mφs, comparing secondary reprogrammed clones (2^o^) to the parental clone (1^o^). Mean ±SD between 3 suid iPSC lines (piPSC2, rhiPSC2 and wbiPSC).

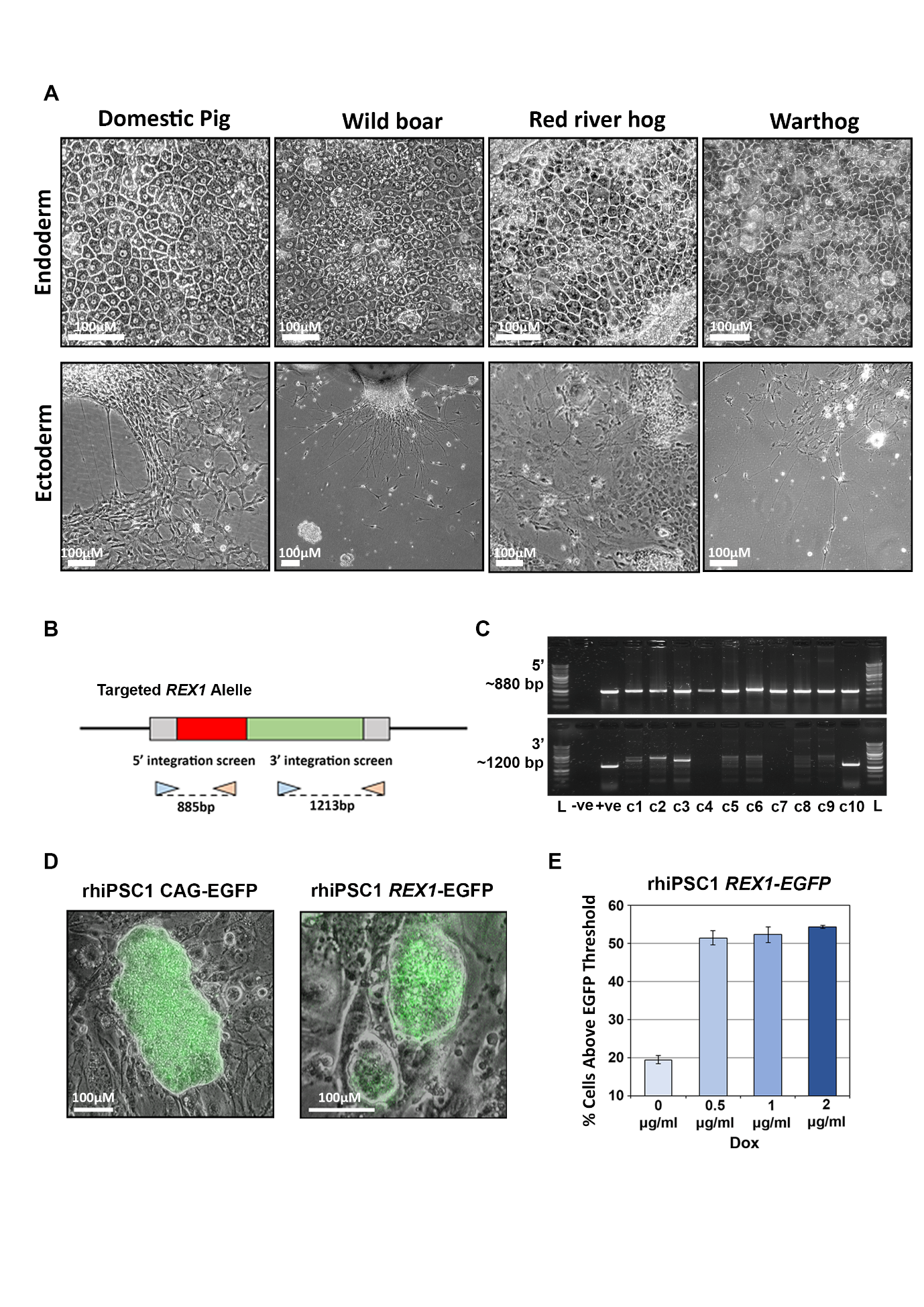

**Figure S4** **Evidence of multilineage differentiation and gene editing potential of suid iPSCs.** (A) Bright field images showing endodermal or ectodermal (neural) morphology in differentiated iPSCs from 4 suid iPSC lines. (B) Diagram showing predicted targeted integration of *REX1-EGFP* construct, and expected product sizes for PCR products identifying correctly integrated construct in rhiPSCs. (C) PCR genotyping of 10 puro-resistant, red river hog EGFP +ve clones. Clones 1, 2, 3 and 10 showed the expected products at both the 5’ and 3’ ends of the integration site. The transfected pool was the positive control (+ve) and wild-type and non-transfected RRH genomic DNA the negative controls (-ve). (D) Composite bright field and fluorescent images showing rhiPSCs expressing CAG-EGFP and targeted REX1-EGFP*.* (E) Bar graph showing a doxycycline (dox) titration in RRH *REX1*-*EGFP* c10. EGFP analysed by flow cytometry at d4 of titration and threshold set to the first replicate in standard 1 µg/mL dox conditions. Mean ±SD of 3 technical replicates.

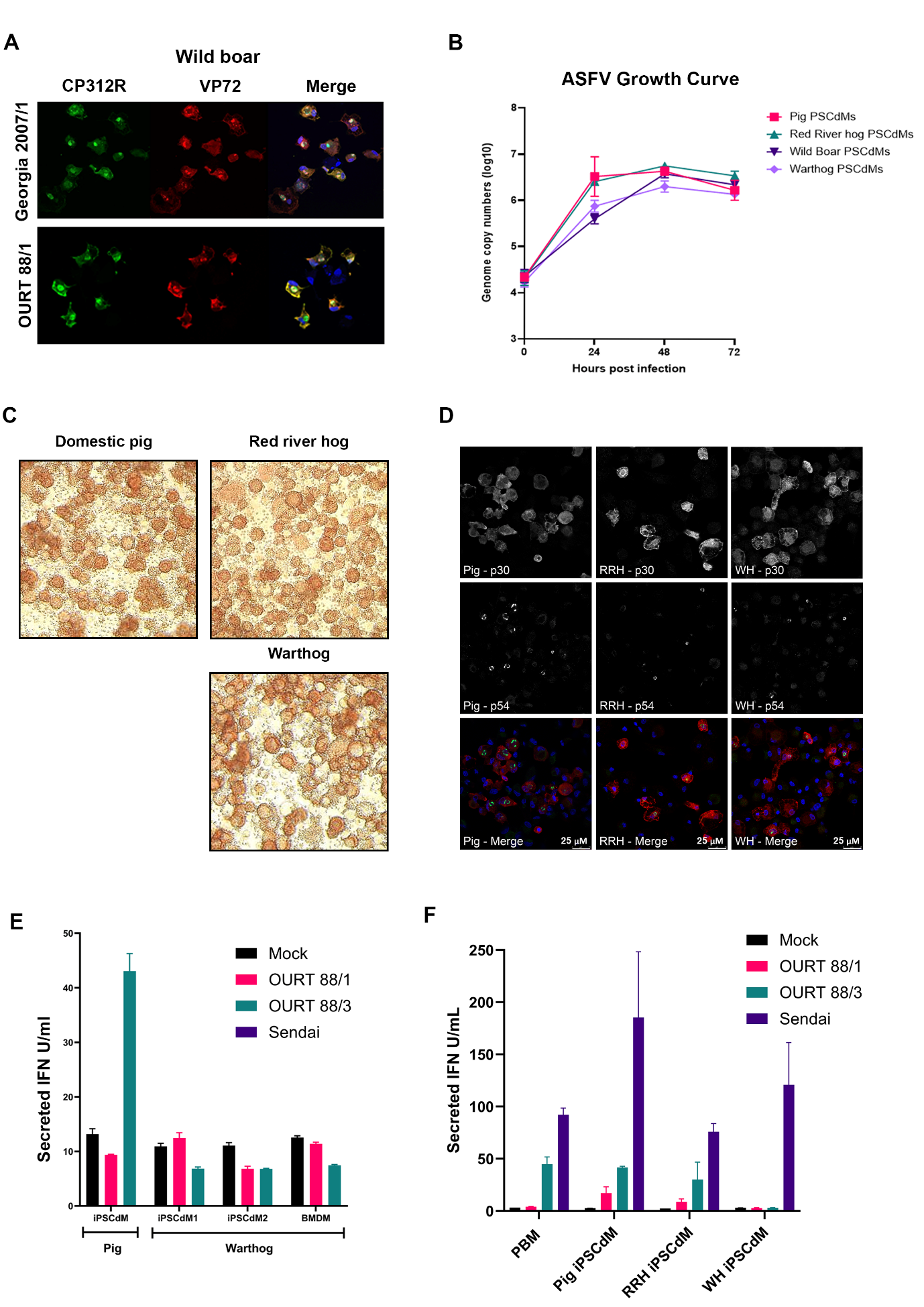

**Figure S5** **Additional characterisation of ASFV infection in ASF relevant domestic and wild pig iPSCdMs** (A) Fluorescent image of ASFV-infected Wild Boar iPSCdMs, 24 hpi, ICC stained for viral proteins CP312R (green) and VP72 (Red). (B) RT-qPCR analysis of ASFV levels (genome copies) present in supernatants from 4 suid iPSCdMs 24 and 48 hpi with ASFV (Georgia 2007/1). Mean and ±SD of 4 independent experiments. (C) Bright-field image showing haemadsorption in ASFV-infected iPSCdMs 24 hpi. (D) Fluorescent images of ASFV-infected (Attenuated OURT 88/3) iPSCdMs from 3 suid species, 24 hpi, stained for viral proteins p54 (green) and p30 (Red). (E) Production of biologically active IFN from piPSCdM1, whiPSCdM1, whiPSCdM2 and *ex vivo* warthog BMDMs after infection with virulent ASFV (OURT 88/1), attenuated ASFV (OURT 88/3) 24hpi. Mean ±SD of 3 independent experiments. Results interpolated using a universal IFNα standard. (F) Data from Figure 4C with addition of Sendai virus +ve for IFN production.

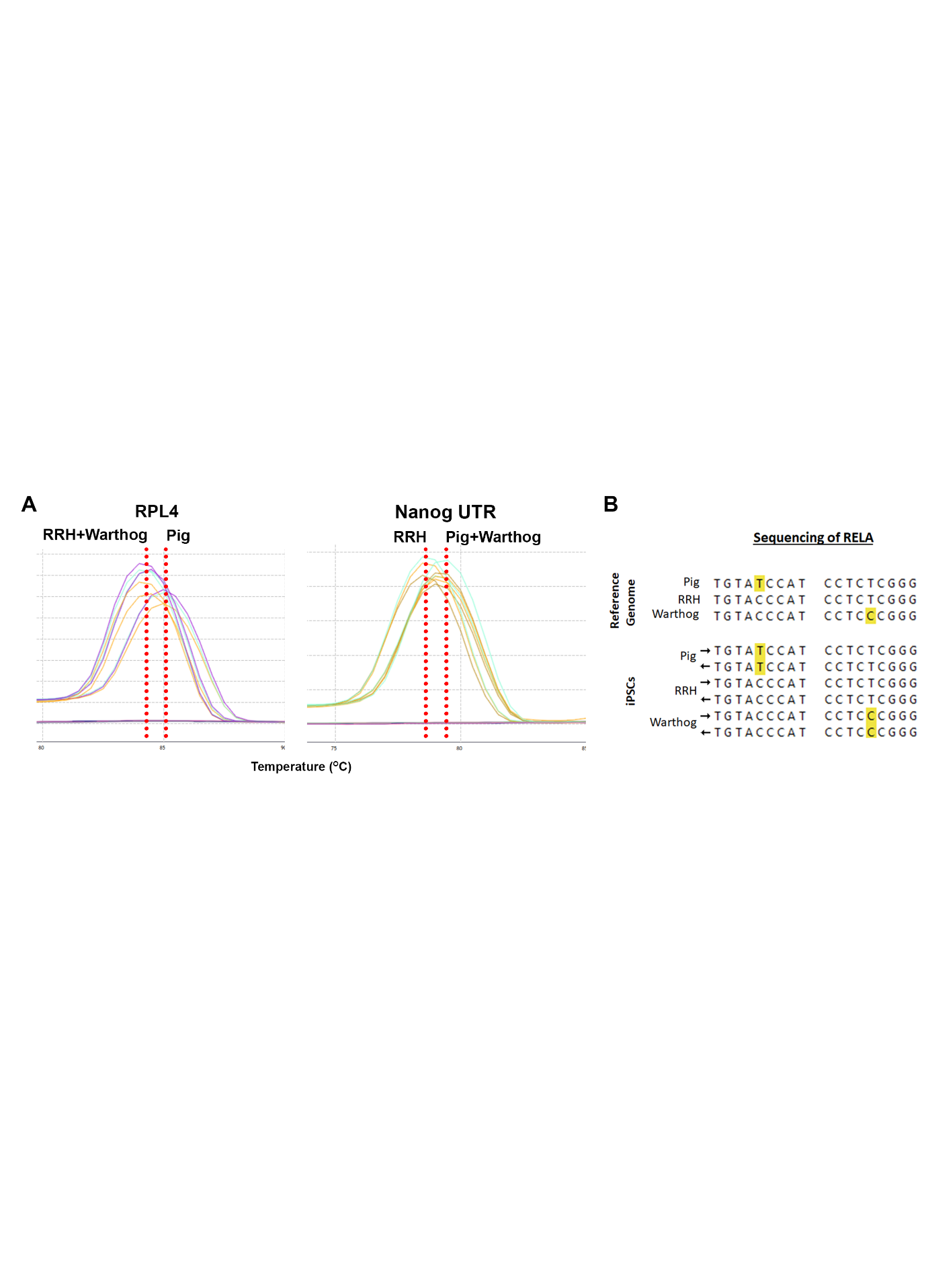

**Figure S6** **Species identification of suid iPSCs** (A) RT-qPCR melt curves using domestic pig specific primers for *RPL4* and *NANOG*. For *RPL4* RRH and warthog have a lower melt curve temperature than domestic pigs. For *NANOG* RRH has a lower melt curve temperature compared to domestic pig and warthog. The combination of the melt curves separates the 3 species. (B) Sequencing of *RELA* in domestic pig, RRH and warthog. Our iPSCs can be identified by previously described SNPs in the *RELA* sequence that match their equivalent reference genome.

**Table S1: qPCR primer list**

| **Gene** | **Sequence** |
| --- | --- |
| ARG1 For | ACTAGGAAGAAAGAAAAGGCCA |
| ARG1 Rev | TTCTTCTGGTGTCTTCCCCA |
| ASFV p72 For | CTG CTC ATG GTA TCA ATC TTA TCG A |
| ASFV p72 Rev | GAT ACC ACA AGA TCA GGC CGT |
| CD163 For | GTGGTCAACTTCGCCTGGTC |
| CD163 Rev | TCAGGTCCCAGCTGTCATCA |
| CD45 For | ACCCTGCTCAGAACGGACAA |
| CD45 Rev | GGGGCACTTGGTGAACTGAC |
| CD86 For | TGCTTGCTAACTTCAGTCAACC |
| CD86 Rev | GCTCAGTGGTTGAATTCTTCGT |
| IFNβ For | GCACTGGCTGGAATGAAACCG |
| IFNβ Rev | AATGGTCATGTCTCCCCTGGGG |
| IL-10 For | TAGGGTGTGCCCTATGGTGT |
| IL-10 Rev | GTGGTTAGGCTTGGAATGGA |
| IL-12β For | TGAGAAGCAGATGTGACCAGA |
| IL-12β Rev | TGGAATCACTTTACAGGAAGCA |
| IL-1β For | TCTGCCCTGTACCCCAACTG |
| IL-1β Rev | CCCAGGAAGACGGGCTTT |
| IL-6 For | ACCGGTCTTGTGGAGTTTCA |
| IL-6 Rev | TTAGGGGTGGTGGCTTTGTC |
| IRF3 For | TTTTCCCGGCTCACTGTACC |
| IRF3 Rev | CACACCCCACTTCTCGTCAG |
| KLF5 For | CCTGAGGACTCATACTGGCG |
| KLF5 Rev | GGTGATCAGAGCGGGAGAAG |
| Nanog For | GAATAGCAACAGTGTGATTCA |
| Nanog Rev | GGAATAGAAGCCCGGGTATT |
| Nanog UTR For | GGTACCCAGCAGCAAATCAT |
| Nanog UTR Rev | TTACGGTGCAGCAGAAATTG |
| NL Plasmid For | TGCACCAGAGTAAGCTGCAC |
| NL Plasmid Rev | CTCCTTTTGATCTGCGCTTC |
| OCT4 UTR For | CAAACTGAGGTGCCTGCCCTTC |
| OCT4 UTR Rev | ATTGAACTTCACCTTCCCTCCAACC |
| OSKM Plasmid For | TCCAAGCAGAGGAGCAAAAG |
| OSKM Plasmid Rev | CCAGCCAATTTCAAGAGAGC |
| PRDM14 For | CCCCAGTGGATGCTTCTCTG |
| PRDM14 Rev | GGTCTTCCTGGGTGAAGTGG |
| Pu.1 For | TACAGGCGTGCAAAATGGAA |
| Pu.1 Rev | AAGTCCCAGTAATGGTCGCT |
| Rex1 For | TCTGAACCCCTCGTGGAAGA |
| Rex1 Rev | AGCTTGCTGTAAGCACCTGT |
| RL Plasmid For | TGCGTGGAGGAAGGAATAAG |
| RL Plasmid Rev | GCAGAGGAAATGGTCAGGTC |
| RPL4 For | AATGTCACTTTGCCTGCTGT |
| RPL4 Rev | CTGGGAATTCGAGCCACAG |
| RPL4 Species ID For | AGGAGGCTGTTCTGCTTCTG |
| RPL4 Species ID Rev | TCCAGGGATGTTTCTGAAGG |
| Sall4 For | CTGCAGATTCATGAGCGCAC |
| Sall4 Rev | GCAATGGTGTTCTCGATGGC |
| SOX2 For | GAGCGCCCTGCAGTACAACT |
| SOX2 Rev | CCCTGCTGCGAGTAGGACAT |
| SOX2 UTR For | ACGGCCATCAACGGTACACT |
| SOX2 UTR Rev | TCTCCTCCCATTTCCCTCTTT |
| TNFα For | GCTCAGCAGTTAGGGACCTG |
| TNFα Rev | AGCCACATCTGCAACCTACC |

**Table S1 qPCR primer list**. A list of all the qPCR primers used in the paper.
